# The pleiotropic functions of Pri smORF peptides synchronise leg development regulators

**DOI:** 10.1101/2023.03.07.531572

**Authors:** Damien Markus, Aurore Pelletier, Muriel Boube, Filip Port, Michael Boutros, François Payre, Benedikt Obermayer, Jennifer Zanet

## Abstract

The last decade witnesses the emergence of the abundant family of smORF peptides, encoded by small ORF (<100 codons), whose biological functions remain largely unexplored. Bioinformatic analyses here identify hundreds of putative smORF peptides expressed in *Drosophila* imaginal leg discs. Thanks to a functional screen in legs, we found smORF peptides involved in morphogenesis, including the pioneer smORF peptides Pri. Since we identified its target Ubr3 in the epidermis and *pri* was known to control leg development through misunderstood mechanisms, we investigated the role of Ubr3 in mediating *pri* function in legs. We found that *pri* play several roles during leg development both in patterning and in cell survival. At larval pupal transition, Pri peptides cooperate with Ubr3 to insure cell survival and leg morphogenesis. Earlier, during larval stage, *pri* activates independently of Ubr3 tarsal transcriptional programs and Notch and EGFR signalling pathways. Our results highlight Ubr3 dependent and independent functions of Pri peptides and their pleiotropy. Moreover, we reveal that the smORF peptide family is a reservoir of overlooked developmental regulators, displaying distinct molecular functions and orchestrating leg development.

**Summary statement:** Pri smORF peptides activate multiple actors regulating signalling pathways, transcription and apoptosis by distinct mechanisms to insure tarsal patterning and epithelial morphogenesis during leg development.

## Introduction

The tremendous development of ribosome profiling, mass spectrometry and bioinformatics revealed the translation of thousands of small Open Reading Frames (smORF, <100 amino acids) in eukaryotes (Wright et al., 2022). As they were considered non-coding due to their small size or their lack of homology, they have been overlooked until recently. SmORF peptides, also known as sORF peptides, microproteins, micropeptides or SEP (sORF encoded-peptides), are translated from smORF located in long non coding (lnc) RNA, or previously alleged lncRNA, in intergenic region or in mRNA, in 5’, 3’UTR or within the coding sequence (Mudge et al., 2022). We are now facing thousands of smORF peptides which require functional characterization to distinguish bioactive smORF peptides from spurious ones. Interestingly, several studies focused on the functions of particular smORF peptides have shown their role in the regulation of different cellular processes involved in development, metabolism or pathologies (Schlesinger and Elsässer, 2022; Wright et al., 2022). For instance, the smORF peptide encoded by Aw112010 lncRNA, highly expressed during infection, has been shown to be required for immunity response (Jackson et al., 2018). Also, Myoregulin and Dworf in mammals, and Sarcolamban in *Drosophila*, all translated from previously annotated lncRNA, control SERCA pump activity in muscles (Anderson et al., 2015; Magny et al., 2013; Nelson et al., 2016). Since smORF peptides have been overlooked so far, they could constitute a reservoir of novel developmental regulators.

The *Drosophila* leg appears to be a good model for testing the biological role of genes encoding smORF peptides because, as an external and segmented organ, the morphology and the possible defects following genetic manipulation of these genes are easily observable in the adult leg. Fly leg development is stereotyped along a proximal-distal axis and relies on the coordination of cell patterning, cell growth, apoptosis and cell morphogenesis (Ruiz-Losada et al., 2018; Suzanne, 2016). Indeed, during embryogenesis, presumptive organs named imaginal leg discs are formed. Then during larval stages, cells proliferate and a complex interplay between morphogens, signalling pathways and transcription factors subdivide the leg disc in different segments separated by folds, that prefigure the future joints. At pupal stage, the leg disc evaginates along the newly formed PD (proximo-distal) axis to form the adult leg composed of seven different segments articulated by joints (Suzanne, 2016).

In *Drosophila*, the pioneer smORF peptides Pri, encoded from a previously alleged lncRNA named *polished rice/tarsal-less* (*pri/tal*), have been firstly identified for their role both in leg formation, more specifically for the development of the tarsus (Galindo et al., 2007), and in embryonic epidermal differentiation (Kondo et al., 2007). The *pri*/*tal* gene is polycistronic and encodes four Pri peptides, which exhibit a conserved motif among arthropods (Galindo et al., 2007; Kondo et al., 2007; Savard et al., 2006). Several studies investigating Pri peptides function during *Drosophila* lifespan have shown they are essential for development or maintenance of various tissues, such as embryonic epidermis and trachea, adult renal and intestinal stem cells and adult legs (Al Hayek et al., 2021; Bohère et al., 2018; Galindo et al., 2007; Kondo et al., 2007). We have previously deciphered their molecular mode of action during epidermal differentiation and showed that Pri peptides interact with the E3 ubiquitin ligase Ubr3 to induce the specific recognition of the transcription factor Shavenbaby (Svb) and ubiquitination of its N-terminal domain. Svb undergoes a ubiquitin-dependent partial degradation of its N-terminal domain, switching it from a large transcriptional repressor (Svb^REP^) form to a shorter activator form (Svb^ACT^), enabling Svb to induce its target genes controlling epidermal differentiation (Kondo et al., 2010; Zanet et al., 2015).

We took advantage of the leg appendage to carry out a functional screen on putative smORF peptides identified specifically in this tissue at two developmental stages corresponding to different disc morphologies. Then, we found that depletion specifically in the leg of 23 of 93 genes encoding for smORF peptides with unknown functions resulted in defects in development. As the most differentially expressed gene at both developmental time points is *pri*/*tal* and their function is not well understood in the developing leg, we decided to investigate its role in the light of our findings (Zanet et al., 2015). Surprisingly, we found distinct functions for Pri during leg development. During pupal stage, the conserved Pri/Ubr3/Svb (Ray et al., 2019) module is involved to ensure cell survival, tarsi morphogenesis and tissue integrity. At the larval stage, Pri peptides are required for EGFR and Notch signalling pathways and transcriptional cascade activation, independently of Ubr3 and Svb. Thus, Pri peptides play pleiotropic functions within the same organ over time by controlling distinct actors, all of which together synchronize morphogenetic events ensuring harmonious leg development.

## Results

### 1- smORF peptides family represents an overlooked reservoir of functional regulators during development

In order to find novel regulatory smORF peptides, we decided to identify candidates and to test their functionality by inducing their loss of function. We focused specifically on the *Drosophila* leg because, as an external organ, it facilitates phenotypic analyses and the identification of defects induced by loss of function of candidate genes. Furthermore, screening in the leg favours linking the type of defects to possible affected signalling pathways implicated for instance in proximo-distal axis patterning, tissue growth, joint formation or epidermal differentiation. To identify genes encoding putative smORF peptides, we performed differential expression analysis combined with a previously published smORF finding approach (Mackowiak et al., 2015). We thus generated transcriptomes of imaginal leg discs at two different stages of development, at wandering larval 3 stage (wL3) before pre-spiracle eversion, which indicates that the peak of ecdysone required for entry into metamorphosis has not yet occurred, and 2 hours APF (After Pupal Formation) (Fig. 1A). Ecdysone signalling induces a transcriptional switch and leg evagination in the proximo-distal axis, possibly favouring our chances to find out regulatory smORF peptides. The bioinformatics analysis to identify smORFs is mainly based on the PhyloCSF method (Lin et al., 2011), which distinguishes coding and non-coding sequences based on substitution patterns in the whole genome alignment of 12 *Drosophila* species. This method allowed us to search for genes encoding for putative smORF peptides (Fig.1A) and to list 396 predicted ones (Table S1), of which 103 are unannotated. Among them, prediction tools identify 162 smORF peptides with specific protein motifs, such as mitochondrial targeting sequence (MitoFates, DeepMito), peptide signal (SignalP 6.0) or transmembrane domain (TMHMM 2.0) (Fig. 1B, Table S1).

**Figure 1.**
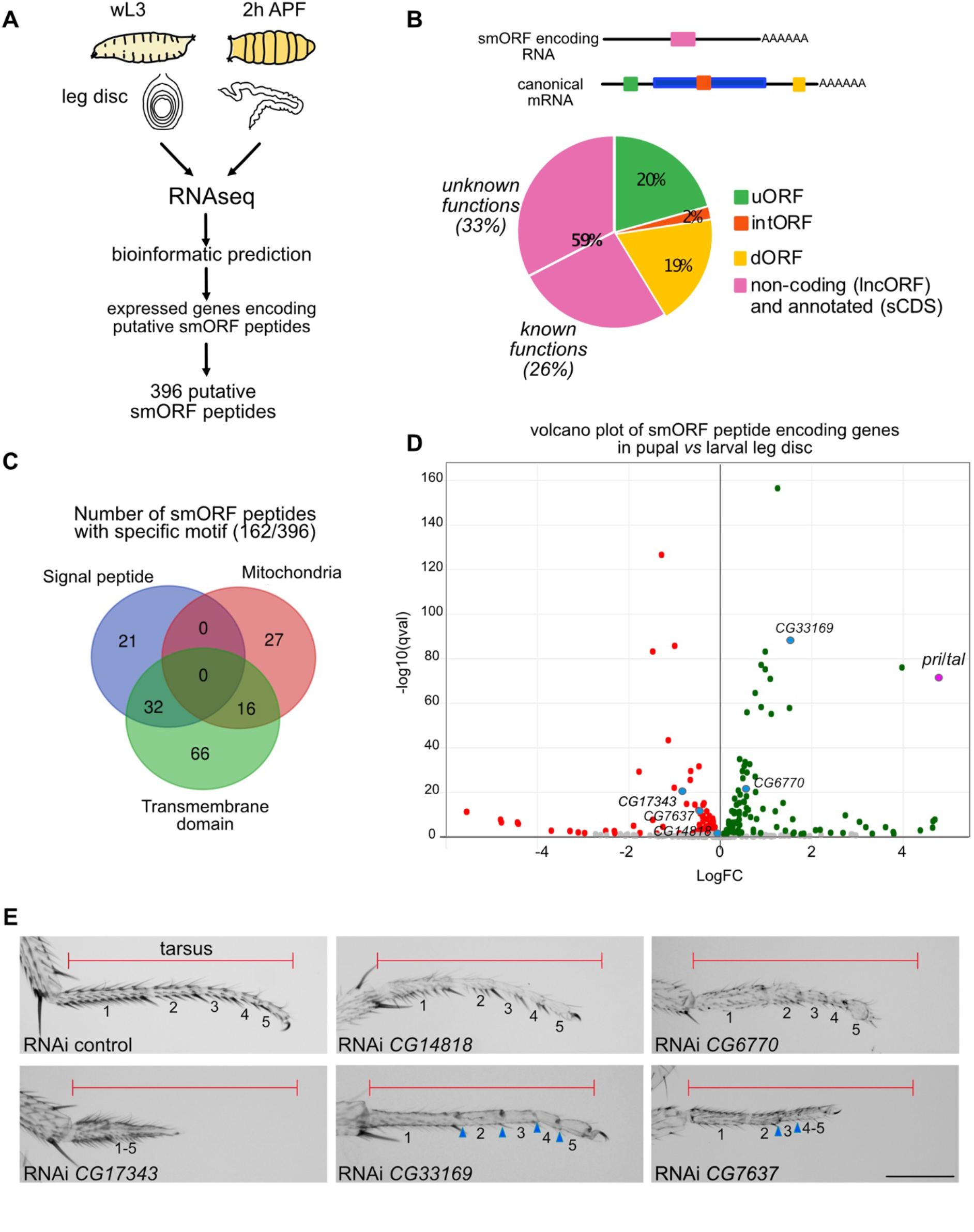
SmORF peptide family comprises a large number of developmental regulators. (A) Schematic representation of the pipeline to identify putative genes encoding smORF peptides expressed in the leg disc. RNAseq was done at two developmental time points of the leg disc, at wandering L3 (wL3) and 2 hours After Pupal Formation (APF). Then bioinformatic analysis was performed to predict smORF peptides from the genes expressed in the leg. (B) Diagram representing the different types of putative smORF peptides found in the leg discs both at wL3 and 2h APF. uORF stands for upstream ORF (green), intORF for internal ORF (orange), dORF for downstream ORF (yellow). SmORF located in monocistronic RNA (pink), which are annotated (sCDS for short coding sequence) or non-coding (lncORF), encoded for either characterised smORF peptides (known functions), or uncharacterised ones (unknown functions). (C) Venn diagram showing predicted protein motif (signal peptides, mitochondria targeting sequence, transmembrane domain) found in 162 putative smORF peptides over the 396 identified. (D) Volcano plot showing log2 fold change values (x-axis) by–log10 corrected p-values (y-axis) for genes encoding for putative smORF peptides between larval and pupal stages. Note that *pri*/*tal* gene is the most differentially expressed gene. (E) Examples of tarsal phenotypes obtained after loss of function of gene encoding smORF peptides. The UAS-RNAi is expressed under the control of *Dll*^EM212^-Gal4 at 29°C. Different defects can be observed, like shortening of the tarsus (compare length of the tarsus with the red line representing the length of the control), fusion of tarsal segments, incomplete joints (blue arrow-heads) and trichome defects. Scale bar = 200μm.

The majority of smORF (59%) are localised in a monocistronic RNA (216 genes), which are either annotated as coding (sCDS for short Coding Sequence) or non-coding (lncRNA and pseudogenes) (Couso and Patraquim, 2017). Remaining smORF are found in canonical coding genes, localised in 5’UTR (uORF), 3’UTR (dORF) or within the main coding sequence (Fig.1C). Of the 216 putative smORF peptides, 96 have been characterised either through conservation among eukaryotes or through functional studies. Then, 120 putative smORF peptides remain with unknown function. Therefore, to figure out which of these smORF are functional and go beyond their theoretical identification, we induced their loss of function during leg development to test their role *in vivo*. Using both the available transgenic fly lines in the stock centres (Bloomington and VDRC) and newly generated fly lines, over these 120 genes, we were able to induce loss of function of 93 of them in the distal part of the leg using the Gal4/UAS system to drive either RNAi or gRNA (CRISPR/Cas9). We used the *Distal-less*-Gal4 driver (*Dll*^EM212^), which is expressed specifically in the distal part of the leg, the tarsus, during *Drosophila* development (Gorfinkiel et al., 1997). Over the 93 tested genes, the depletion of 23 of them impaired tarsus formation. We observed different types of tarsal defects, like fusion of segments, incomplete joints, necrosis, tarsi reduced size, altered cuticle formation or trichome pattern (Fig. 1E and Fig. S1). The diversity of phenotypes suggests that smORF peptides are implicated in different cellular processes. Notably, loss of function of a high proportion of tested genes (24%), which encode putative smORF peptide, induces phenotypes. Therefore, our functional screen highlights the smORF peptides as a reservoir of novel developmental and cellular regulators.

Interestingly, differential expression analysis at two developmental time points shows a remarkable switch in gene expression between larval and pupal stage (946 genes with log2FC>1 and 827 with log2FC<1). Among the genes encoding putative smORF peptides, the most differentially expressed gene is the *pri*/*tal* gene (log2FC=4,77; Fig. 1D, Table S1). Pri peptides have been discovered for their role in tarsus formation (Galindo et al., 2007), where they are known to control tarsal patterning and joint formation (Natori et al., 2012; Pueyo and Couso, 2008; Pueyo and Couso, 2011). Pri peptides are required for the establishment of the transcriptional program controlling tarsal segmentation, but the underlying mechanisms are not known. Also, it has been proposed that Pri peptides control joint morphogenesis through Svb and Notch regulation during pupal stage (Pueyo and Couso, 2011). As the molecular mechanisms of action of Pri peptides remain not well understood during larval and pupal leg development, we then investigated their functions in the light of our recent findings and analysed the role of Ubr3 (Zanet et al., 2015).

### 2- Pri peptides play distinct roles at larval and pupal stages

As we observed a strong increase in *pri* expression between wL3 stage and 2h APF stages (Fig. 1D), we analysed *pri* mRNA localisation by quantitative fluorescent *in situ* hybridization (smiFISH) (Couturier et al., 2019). As previously described (Galindo et al., 2007), we observed *pri* mRNA at midL3 stage in the form of a ring-shaped pattern marking the presumptive territory of the tarsus, which stops at wL3 stage (except in the notochordal organ). Then, at the onset of metamorphosis, *pri* is strongly reactivated in the whole leg disc and in the peripodial membrane (Fig. 2A). The dynamic pattern of *pri* expression during leg development suggests different functions. To test this hypothesis, *pri* expression was specifically depleted in the tarsus at larval or/and pupal stages by using different genetic approaches (Fig. 2B). To analyse the effect of *pri* depletion only during the larval stage, we used the *tal*^1^ mutant (Galindo et al., 2007), in which *pri* expression is specifically absent during the larval stage in the leg, but unaffected at the onset of metamorphosis (Fig. 2C). Indeed, in our hands, depletion of *pri* with RNAi during the larval stage was not efficient enough to get rid of the larval function of *pri. tal*^1^ allele affects the *cis*-regulatory genomic region controlling larval *pri* expression in the leg, named *pri*I (Dib et al., 2021). Indeed, ectopic expression of *pri* under the control of *pri*I in *tal*^1^ mutant background restores tarsus morphology (Fig. S2A). Thus, the absence of *pri* specifically during the larval stage leads to the fusion of the tarsal segments and then to a shorter tarsus (Fig. 2B). To specifically delete *pri* at pupal stage, we used the Gal4/UAS and the thermo-inducible systems (*Dll*>RNAi *pri*; *tub*Gal80ts). The absence of *pri* during pupal stage induces the loss of tissue integrity of the distal part of the leg (Fig. 2B). Then, to perform *pri* depletion during both larval and pupal stages, we induced large *pri*^-/-^ clones in *Minute* context specifically in the tarsus, using the *FRT*/*FLP* system, in which the *flippase* is expressed under the control of *Dll*. Continuous depletion of *pri* over larval and pupal stages accumulates both phenotypes and results in a shorter and dramatically altered tarsus (Fig. 2B). Therefore, during leg development, Pri smORF peptides exhibit distinct functions as they are required for patterning proximo-distal axis in the leg disc and consequently tarsal segmentation, and then at pupal stage, they are essential to ensure tissue integrity.

**Figure 2.**
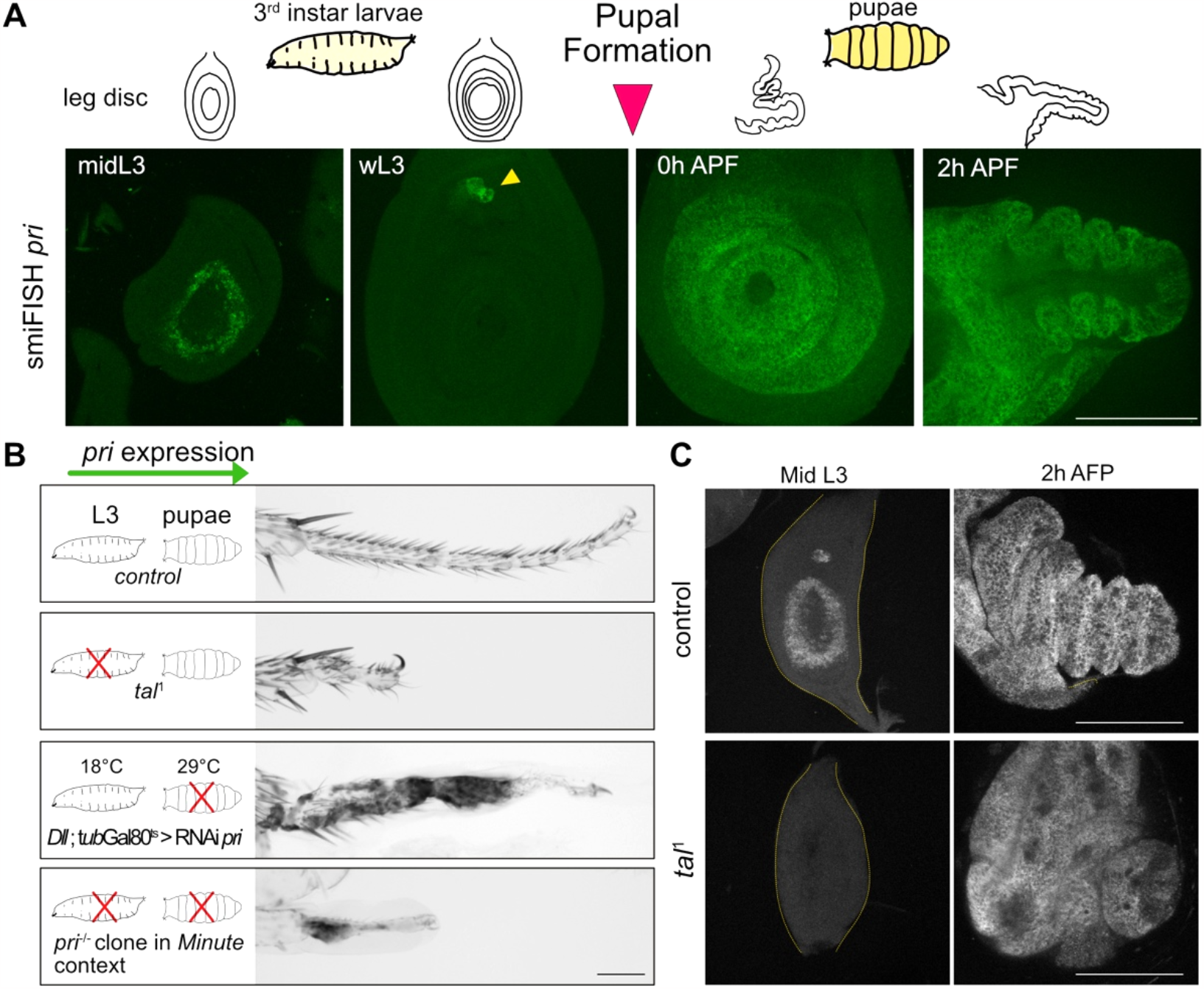
Pri peptides have distinct roles during leg development. (A) Fluorescent *in situ* hybridisation of *pri* mRNA at different stages of leg disc development, schematised above by the drawings: from left to right, midL3 for mid third instar larvae 3, wL3 for wandering third instar larvae 3, 0h APF (After Pupal Formation) corresponds to the start of metamorphosis, 2h APF. *pri* expression is transient and pulsatile in a typical ring-shaped pattern in the presumptive tarsal region in the midL3 disc and *pri* expression is abolished in the wL3 disc (except *pri* expression observed in the chordotonal organ, yellow arrow-head). At the metamorphosis, corresponding to 0h APF, *pri* is strongly expressed in the whole distal region. In 2h APF disc, *pri* is expressed in the epithelium of the disc and in the peripodial membrane. (B) Phenotypes of adult leg obtained after stage specific *pri* depletion, schematised by a red cross, during larval or/and pupal stages. *pri* depletion during larval stage, here obtained by using the *tal*^*1*^ mutant, induces a strong reduction in tarsal size, fusion of the different tarsal segments and absence of tarsal joints. Loss of function specifically during pupal stage is induced by expressing UAS-RNAi *pri* under the control of *Dll*^EM212^-Gal4 driver and *tub*-Gal80^ts^ at 29°C, and leads to tissue integrity loss. To deplete *pri* during larval and pupal stages, *pri*^-/-^ (*tal*^S18^) clones in the *Minute* cellular context were induced specifically in the tarsus (*Dll*^EM212^>Flippase). The resulting phenotype cumulates the defects described above, up to the tarsus disappearance. (C) *pri in situ* fluorescent hybridisation (smiFISH) in control and in *tal*^*1*^ leg discs. *pri* expression is absent in the *tal*^*1*^ mutant specifically during larval stage (midL3, outlined by the yellow dashed-line) and reactivated at metamorphosis (2h APF). Note the persistence of fusion of the tarsal segments in *tal*^*1*^ disc at 2h APF. All scale bars = 100μm.

### 3- Pri peptides are required for early steps of tarsal patterning

During larval stage, tarsal patterning is regulated by multiple actors, that define each segment composing the future tarsus. During the first and the second instar larvae, the morphogens Hedgehog (Hh), Decapentaplegic (Dpp) and Wingless (Wg) establish the anterior-posterior and dorso-ventral axis in the leg disc (Lecuit and Cohen, 1997). Consequently, the expression of *dll* is activated in the leg disc, then defining the tarsal region during L3 stage (Diaz-Benjumea and Cohen, 1994). Also, during early L3 stage, the EGFR signalling pathway is activated through the integration of the signal of Wg, Dpp and Dll at the center of the disc, known as the EGFR organizing center (EOC), and will govern the identity of the pretarsus. At mid-late L3 stage, a second wave of EGFR signalling (non-EOC), mostly dependent on the metalloprotease Rhomboid and the ligand Spitz, is activated in the tarsal region (Campbell, 2002; Galindo et al., 2002; Newcomb et al., 2018). Thus, both EGFR and Dll subdivide the medial tarsal region and allow the expression of *spineless* (*ss*), which in turn induces *rotund* (*rn*), both TFs being necessary for subsequent tarsal patterning (Emmons et al., 2007; Natori et al., 2012). Furthermore, Notch signalling is required for patterning boundaries between segments, which prefigure joint formation (de Celis et al., 1998). Notably, *rn* is necessary for Notch pathway activation (St Pierre et al., 2002). During tarsal patterning, it was previously shown in *tal*^1^ and *tal*^KG^ mutants, or in *tal*^S18^ mutant clones, that *pri* was required for activating *ss* and *rn* transcription (Galindo et al., 2007; Natori et al., 2012; Pueyo and Couso, 2008). Here we showed in the *tal*^1^ mutant, in which *pri* expression is specifically abrogated in leg disc at larval stage, that Dll protein is still present while Ss and Rn are absent (Fig. 3A, Fig. S3A). We then investigated at which stage of the regulatory cascade *pri* was acting. We observed that Notch signalling pathway, which is activated from the larval stage, was absent in the presumptive region of the tarsus in *tal*^1^ (Fig. 3B), as confirmed by the absence of Deadpan (Dpn) protein and *dysfusion*-lacZ reporter line activity, both direct targets of the Notch pathway (Córdoba and Estella, 2014). As EGFR is important for limiting Notch signalling at joint boundaries (Shirai et al., 2007), we stained *tal*^1^ L3 leg discs with anti-Phospho-ERK antibody, a marker of MAPK activity used as a read-out of active EGFR signalling pathway, and revealed its absence in the tarsal region (Fig. 3C). Furthermore, we found that in *tal*^1^ *rhomboid* mRNA was absent in tarsal region, showing that the second wave of EGFR activation is compromised (Fig. 3D). Nevertheless, in the absence of *pri*, the initial EGFR wave is activated as indicated by the presence of the TF Clawless specific from the pretarsus and the formation of the claws (Fig. S3B) (Kojima et al., 2005). Therefore, our data reveals that Pri peptides are required for *rhomboid* transcription, and consequently EGFR signalling, Notch signalling and activation of the tarsal transcriptional program.

**Figure 3.**
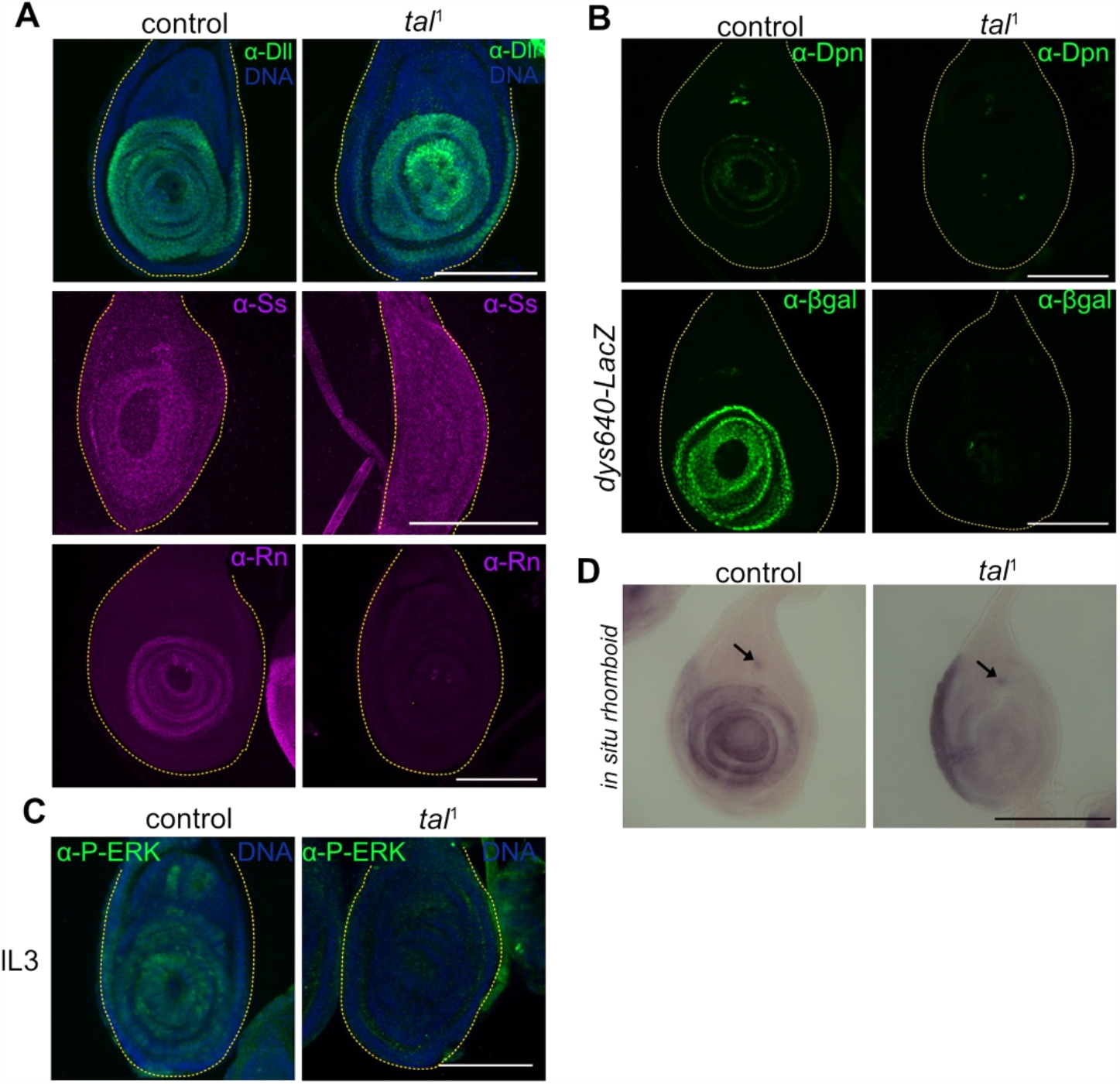
Pri peptides control tarsal patterning by activating the transcriptional program and the Notch and EGFR signalling pathways. (A) Immunostaining of Dll, Ss and Rn transcription factors in control and in *tal*^1^ mutant discs at L3 stage. Dll is present in both control and *tal*^1^ distal region, whereas Ss and Rn proteins are absent in the *tal*^*1*^ mutant leg discs. (B) Immunostainings of Deadpan (Dpn) and βGalactosidase (βGal) in a control and *tal*^*1*^ in L3 leg disc. *dys640*-LacZ is an enhancer of *dysfusion* gene directly activated by Notch, showing the absence of Notch signalling in the tarsal region. (C) The anti-P-ERK signal, used as a read-out of EGFR signalling, stains at late L3 stage (lL3) the whole tarsal region in the control while being absent in the *tal*^*1*^ mutant. The leg discs are outlined by the yellow dashed-line. *In situ* hybridisation of *rhomboid* mRNA in control and in *tal*^1^ leg discs. (D) The *rhomboid* mRNA is observed in the presumptive tarsal region in concentric ring pattern. In the *tal*^*1*^ mutant, *rhomboid* expression is abrogated in the disc except in the chordotonal organ (black arrow). All scale bars = 100μm.

We conducted genetic epistasis analysis to determine the functional order of these genes in tarsal patterning. Interestingly, ectopic expression in the tarsus of *tal*1 mutant of activated forms of EGFR, sSpitz, *ss* or *rn* failed to restore Notch and EGFR signalling, or transcription factors activation (Fig. S3D). Ectopic expression of *ss* is not sufficient to activate *rn*, and ectopic expression of *rn* is not sufficient to activate Notch signalling (Fig. S3C,D). None of these actors is able to replace *pri* function, suggesting that Pri peptides are acting at several steps in this molecular cascade. Furthermore, ectopic expression of *pri* in the *wg* domain, which is expressed in a sub-region of the leg during development (Diaz-Benjumea and Cohen, 1994), enables Notch signalling pathway and tarsal transcriptional program activation, as visualised respectively with Dpn and Rn immunostainings (Fig. 4A). Also, these factors were reactivated beyond the area of *wg* expression domain, over a distance of several cell diameters (Fig. 4B), illustrating the non-cell autonomous properties of Pri peptides, which were previously observed (Galindo et al., 2007; Kondo et al., 2007; Chanut-Delalande et al., 2014). We noted that the level of Dpn and Rn proteins were similar within and out the domain of *wg* expression (Fig. 4B), suggesting that activation is not Pri dose-dependent.

**Figure 4.**
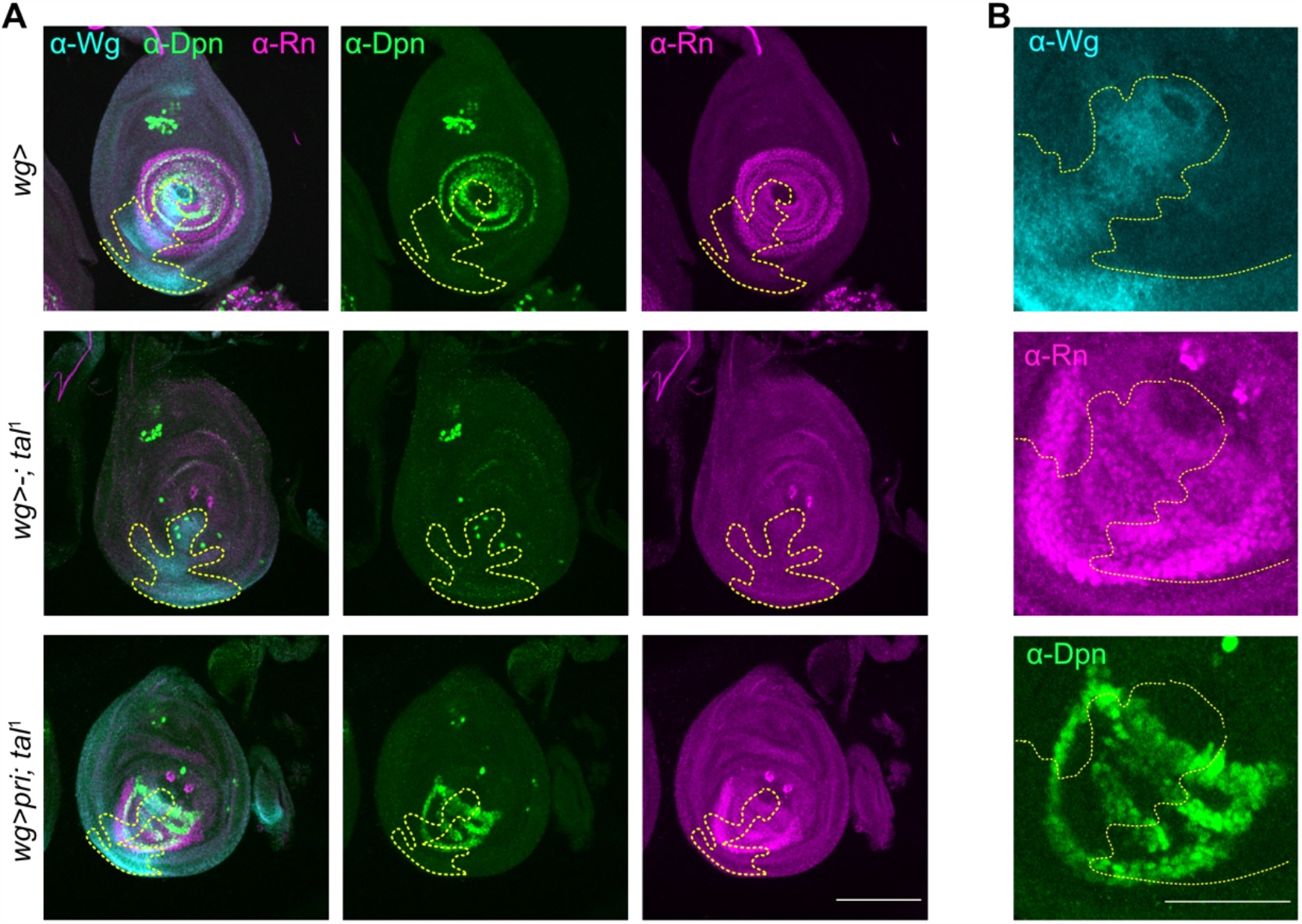
Cell non-autonomous properties of Pri peptides synchronise transcriptional program and the Notch and EGFR signalling pathways in tarsus. (A) Rescue experiments have been conducted by expressing ectopically *pri* in *tal*^1^ mutant background under the *wingless*-Gal4 driver (*wg*). Wg staining delimits the region where *pri* is expressed (outlined by the yellow dashed-line). Rn and Dpn are used as read-out of tarsal patterning and Notch signalling pathway activations. Rn and Dpn are normally patterned in the control disc, whereas they are absent in the *tal*^1^ mutant. Ectopic expression of *pri* (*wg*>*pri*; *tal*^*1*^) rescues partially the *tal*^1^ phenotype since Rn and Dpn stainings are restored in the *wg* region. Scale bar = 100μm. (B) Magnification on the *wg* region of the *wg*>*pri*; *tal*^*1*^ leg disc. Note that Rn and Dpn are present at the same level far beyond the *wg* region, showing that Pri peptides induce cell non-autonomously activation of tarsal patterning. Scale bar= 50μm.

In conclusion, our data show that Pri peptides are required for the activation of signalling pathways and transcription factors that governs tarsal patterning. Non-cell autonomous properties of Pri peptides may coordinate the activation of these actors within the tarsus to ensure harmonious tarsal development.

### 4- Pri peptides induce Svb processing in Ubr3 dependent manner at larval-pupal transition

To go further, we then investigated whether the roles of Pri peptides during leg development were dependent on Svb and Ubr3, the partners identified for mediating Pri functions for trichome formation during epidermal differentiation (Kondo et al., 2010; Zanet et al., 2015). Both *Ubr3* and *svb* are expressed during larval and pupal stages at comparable levels (Fig. S2B). The depletion of *svb* and *Ubr3* by RNAi under the control of *Dll*-Gal4 driver induces strong defects in tarsus morphology, showing that both proteins are required for leg development (Fig. S2C). The Pri peptides and Ubr3 induce Svb processing, switching the Svb^REP^ to the Svb^ACT^ forms (Zanet et al., 2015). We generated a fly line in which the endogenous Svb was tagged to the GFP at the C-terminal position (KI *svb*::*GFP*, Fig. 5A). Therefore, immunostainings against both GFP protein and 1S domain, which is specific from the Svb^REP^ (Kondo et al., 2010), allow to visualize Svb^REP^ form (1S and GFP positive) and Svb^ACT^ form (only GFP positive) during leg disc development. We revealed that Svb is ubiquitously expressed in the imaginal disc of the leg during development, since it is present as the repressor form in the larval stage and as the activator form in the pupal stage (Fig. 5B, Fig. S2D). Thus, Svb processing is occurring at larval-pupal transition, at the time of tarsus eversion. We further found that Svb processing relies on Ubr3 and Pri peptides as shown by the persistence of the 1S signal at pupal stage when any of these genes was depleted by clonal analysis (Fig. 5C). Therefore, the function of Pri/Ubr3/Svb module is reiterated during development, specifically in the imaginal leg discs during the larval-pupal transition when *pri* expression is strongly reactivated.

**Figure 5.**
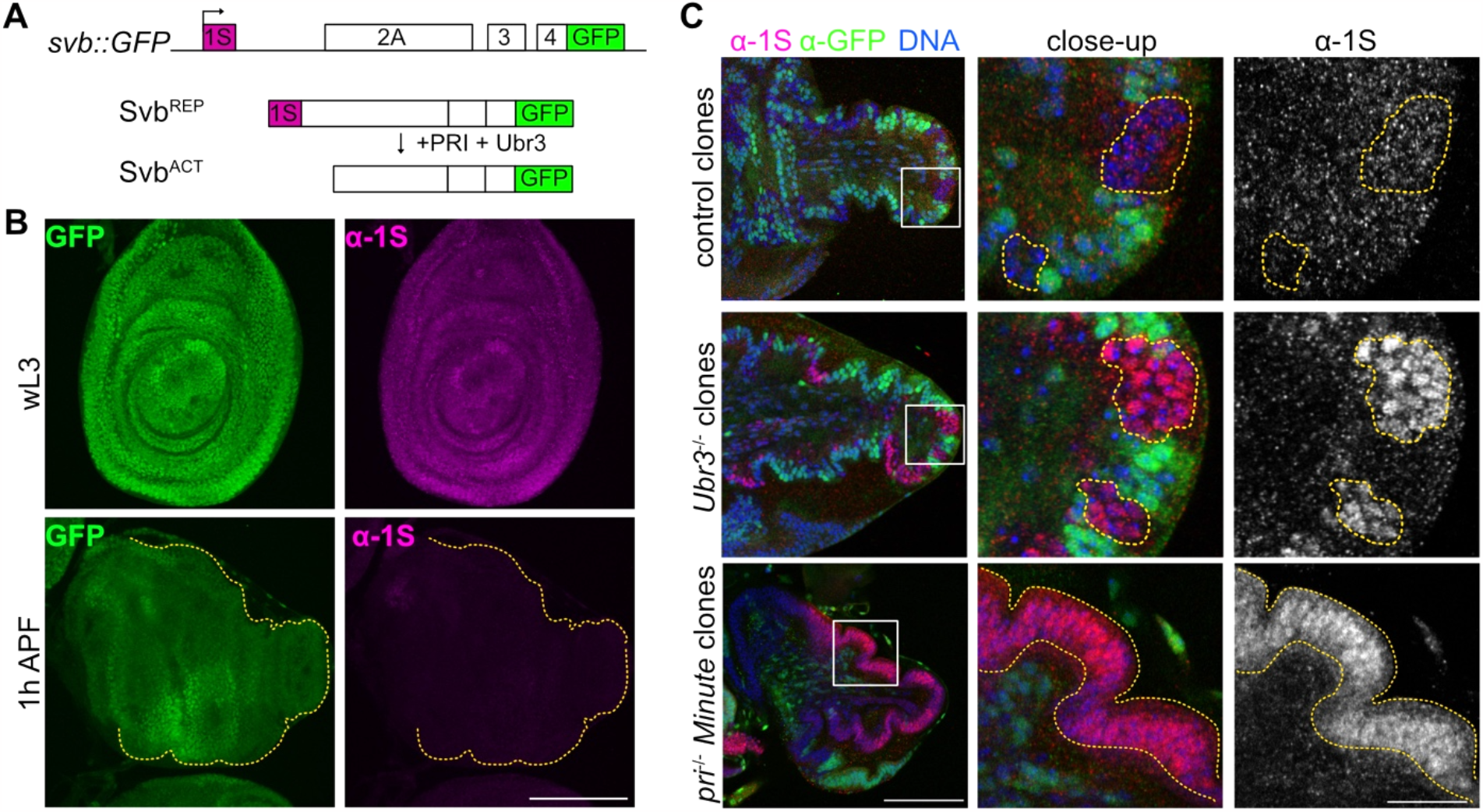
*Pri* and *Ubr3* are required for Svb processing at the larval-pupal transition. (A) Drawing representing the knock-in (KI) of GFP in the *svb* locus (*svb*::*GFP*). *svb* is transcribed and translated as a long repressor form (Svb^REP^), that contains the exon 1S. In embryonic epidermis, the presence of Pri peptides and Ubr3 induce the N-terminal domain degradation, leading to a shorter activator form of Svb (Svb^ACT^). (B) Anti-GFP (green) and anti-1S (purple) immunostainings on *svb*::*GFP* KI at larval stage (wL3) and pupal leg disc (1h APF, contoured by yellow dashed-line) showed that Svb is localised ubiquitously in the leg disc and is under Svb^REP^ form. However, at the pupal stage, Svb is processed in the whole leg disc, and remains as the Svb^ACT^ form, as confirmed by the absence of 1S signal. Scale bar = 100μm. (C) Anti-1S immunostaining in the pupal leg disc where mosaic clones for *pri*^-/-^ (*tal*^S18^) and *Ubr3*^/-^ (*Ubr3*^B^) were induced. The clones are indicated by the absence of GFP and contoured by the yellow dashed-line. The DNA is marked in blue, 1S in red and the GFP in green, white square highlights the region displayed in the close-up. In control clones, Svb is processed as 1S signal is lost. In *Ubr3*^-/-^ clones, 1S staining is remaining, showing that Svb^REP^ is not processed. p*ri*^-/-^ mutant clones have been generated in *Minute* background with the flippase under the control of *Dll*^EM212^-Gal4 in order to get large clones (if *pri*^-/-^ clones are too small, they are behaving like control clones). The absence of GFP indicates that almost all the leg is clonal (yellow dashed-line) and positive for anti-1S signal. Scale bar= 100μm. On the close-up, scale bar= 20μm.

### 5- Pri peptides control cell survival during pupal stages

Our results showed that the Pri/Ubr3/Svb module is required for pupal leg development. Interestingly, depletion of each of these genes induces distinct phenotypes in terms of severity, suggesting that Pri peptides possess additional developmental functions compared to Ubr3 and Svb. Indeed, RNAi depletion of *pri* leads to a more severe phenotype than the depletion of *Ubr3*, which in turn is more severe than *svb* depletion (Fig. 2C, Fig. S2C). As the absence of *pri* during pupal stage induces the loss of tissue integrity, we tested whether cell death could be the cause of this phenotype. We thus stained leg disc with anti-Dcp-1, the cleaved form of the ortholog of human caspase-3, and we observed a strong increase in the number of apoptotic cells (Fig. 6A), thus corroborating a role for Pri peptides in protecting cells from apoptosis. Then, we induced loss of function of *Ubr3* and *Svb* to test their role in cell survival. Generation of *Ubr3*^-/-^ clones in imaginal leg disc induced an increase in apoptosis (Fig. 6B), showing that Ubr3 is required to protect cells from death. Although variable, *svb* depletion induces significantly more Dcp-1 positive cells, indicating that Svb also plays a role in protecting cells from cell death (Fig. S4A,B).

**Figure 6.**
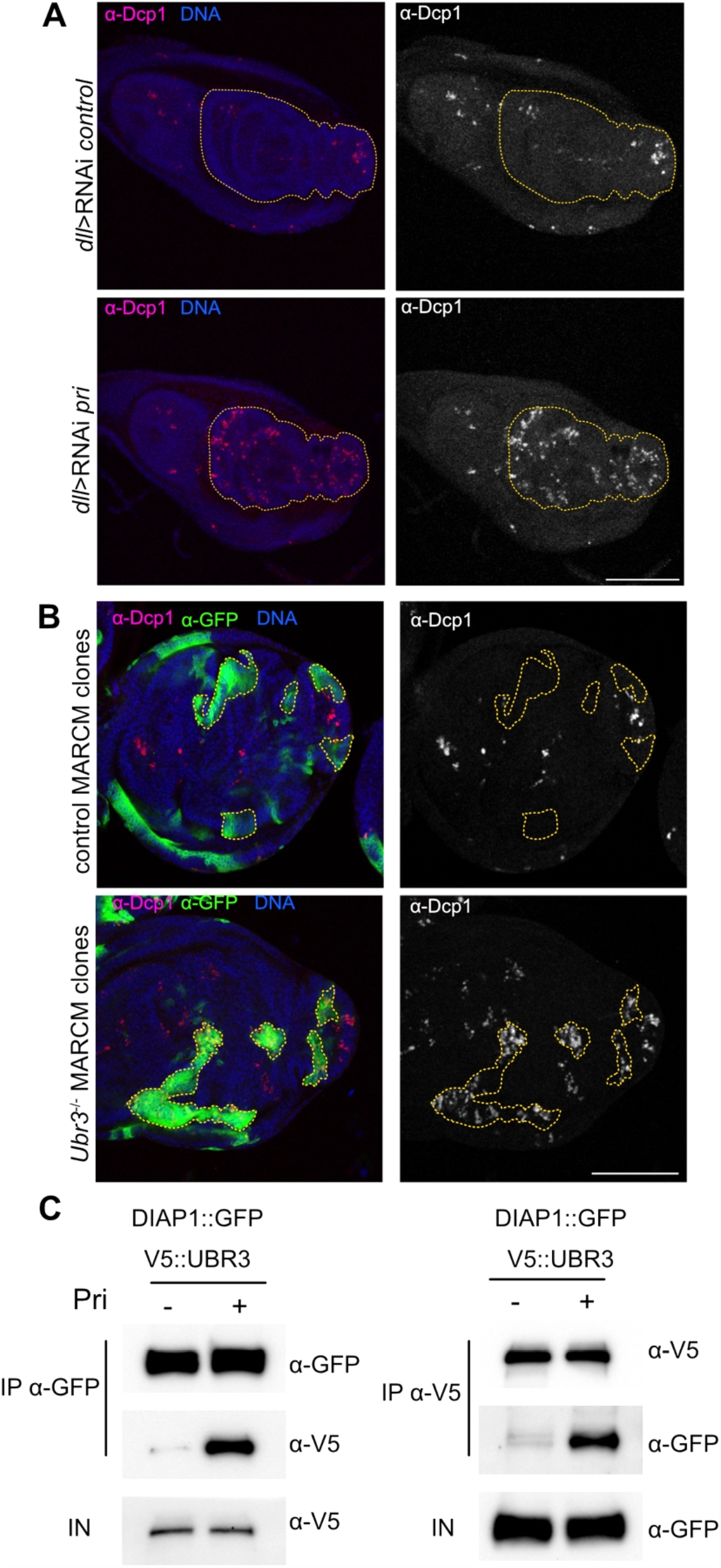
Pri peptides and Ubr3 protect cells from apoptosis. (A) Anti-Dcp-1 antibody stains apoptotic cells (in red) in larval leg disc, that are present in the control leg disc (*Dll*^EM212^>RNAi *luciferase*), specifically in the pretarsus, as often described. Depletion of *pri* by expressing UAS-RNAi *pri* under the control of *Dll*^EM212^-Gal4 driver induces a dramatic increase in the number of apoptotic cells. Scale bar= 100μm. (B) Anti-Dcp-1 staining of leg disc where control and *Ubr3*^-/-^ MARCM clones are induced (the clones are GFP positive, in green). We observe a strong increase of Dcp-1 positive cells in *Ubr3*^-/-^ MARCM clones. Scale bar= 100μm. (C) Co-immunoprecipitation of Ubr3::V5 and DIAP1::GFP with or without Pri peptides. DIAP1::GFP and Ubr3::V5 are co-expressed in S2 cells with or without *pri*, and then co-immunoprecipitated with anti-GFP (left panel) or anti-V5 (right panel) antibodies. In the absence of *pri*, DIAP1 and Ubr3 co-interact barely. With Pri peptides, this interaction strongly increases.

To go further, we investigated the role of the module Pri/Ubr3/Svb in the regulation of the major anti-apoptotic factor DIAP1. Since Svb was shown to protect cells from apoptosis via the regulation of *DIAP1* transcription in digestive stem cells (Al Hayek et al., 2021; Bohère et al., 2018), we performed *DIAP1* fluorescent *in situ* hybrization (smiFISH) in imaginal leg discs in the absence of *svb*. We did not observe change in the level of mRNA *DIAP-1* during the time window encompassing larval-pupal transition, suggesting an alternative mechanism by which Svb protects cells from apoptosis (Fig. S4C). Interestingly, Ubr3 was shown to protect cells from apoptosis in imaginal eye disc through its interaction and protection of DIAP1 protein (Huang et al., 2014), we thus tested the influence of Pri peptides on Ubr3/DIAP1 protein interaction in *Drosophila* S2 cells. Even though weak co-immunoprecipitation between Ubr3 and DIAP1 is observed without Pri peptides, the presence of Pri increases massively the interaction (Fig. 6C). Our data suggests that Pri peptides and Ubr3 are cooperating for protecting DIAP1 from degradation and prevent cells from entering in apoptosis *in vivo*.

Altogether, our data reveals that the module Pri/Ubr3/Svb is protecting the leg from cell death and is necessary for morphogenesis and preserving tissue integrity throughout the development of the pupa.

### 6- Pri controls larval disc patterning in Svb/Ubr3 independent manner

As we found that the module Pri/Ubr3/Svb is important during pupal development, we then challenged the role of Ubr3 and/or Svb in mediating Pri peptides functions during larval stages. First, we generated *svb* loss of function by generating mutant clones or inducing RNAi and stained leg discs to analyse the effects on tarsal patterning. Persistence of Ss and Rn TFs or Notch signalling pathway in the absence of *svb* demonstrate that Svb is not required for their activation (Fig. 7A, Fig. S5A). Strikingly, Svb, which is under its full-length repressive form at this stage of development, is fully degraded at the timing of transient *pri* expression (Fig. S5B). This complete degradation is dependent of *pri* because in *tal*^1^ mutant, Svb protein persists (Fig. S5B). Then, we depleted *Ubr3* by the same genetic approaches, either by generating *Ubr3* mutant clones or by inducing loss of function by RNAi. Notch pathway is not affected by loss of *Ubr3*, as visualised by the persistence of Dpn in *Ubr3* mutant clones (Fig. S5A). Additionally, Ss and Rn are still present in the *Ubr3*^−/ −^ clones, even though the signal is ranging from an unaffected to lower protein level (Fig. 7B). Since the absence of *Ubr3* leads to apoptosis, the variation on Ss and Rn protein levels might be the consequence of a deleterious cellular context. Depletion of *Ubr3* by RNAi resulted to the same observation, supporting that Ubr3 is not required for mediating *pri* function in the larval leg disc (Fig. 7C).

**Figure 7.**
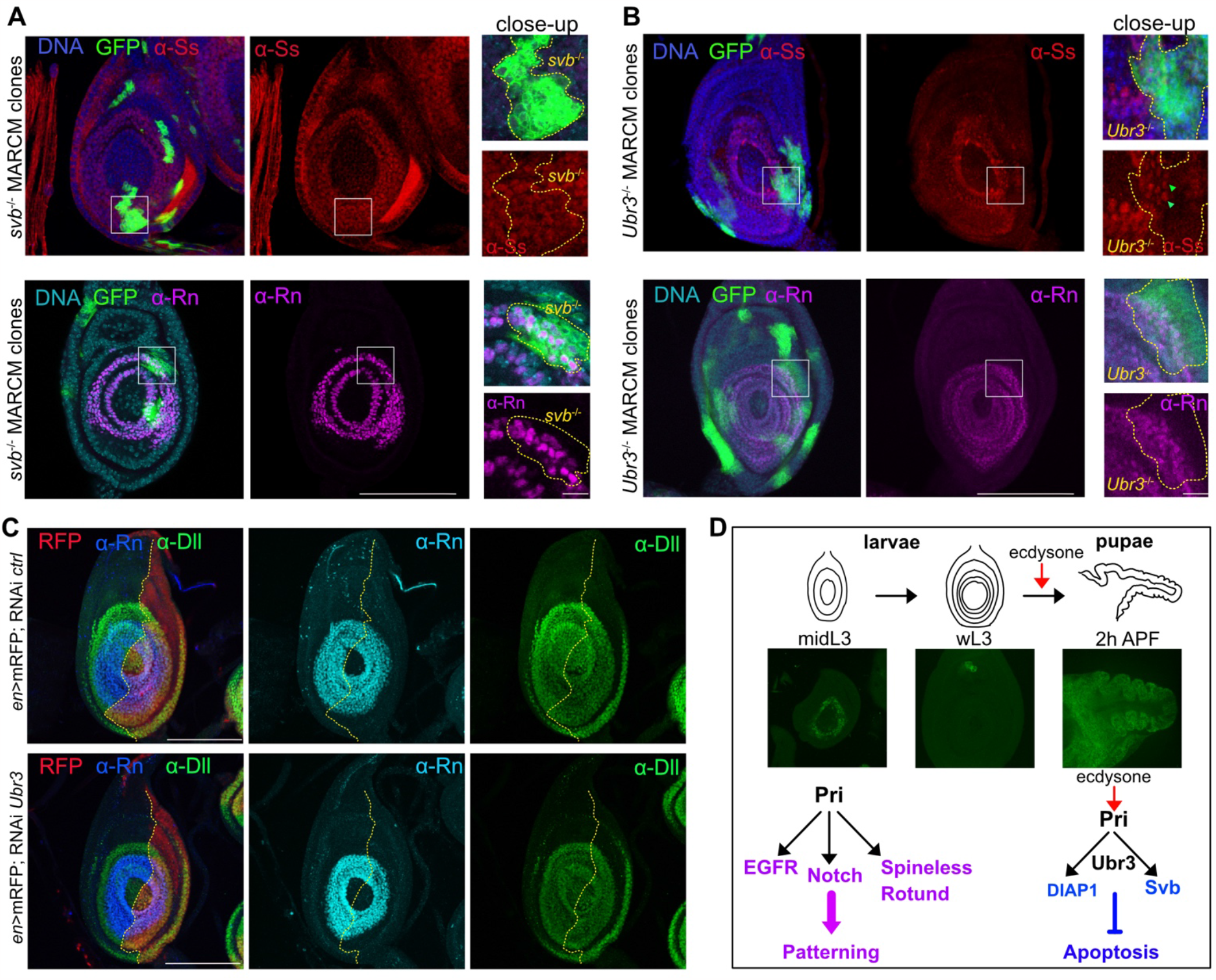
Pri peptides activate tarsal TF and EGFR and Notch signalling pathways in Svb/Ubr3 independent manner. (A) Ss and Rn were stained in *svb*^-/-^ (*svb*^PL107^) MARCM clones in L3 leg discs. Clones are in green and outlined in yellow in the close-up. In *svb*^-/-^ clones, both Ss and Rn proteins remain. (B) Ss and Rn were stained in *Ubr3*^-/-^ (*Ubr3*^B^) MARCM clones in L3 leg discs. In the *Ubr3*^-/-^ clones, both Ss and Rn proteins remain, though the level of Ss is lower (green arrow-heads). (A,B) White square highlights the region displayed in the close-up showed on the right of the panel. Scale bar= 100μm, scale in the close-up= 10μm. (C) UAS-RNAi *luciferase* (*ctrl*) or UAS-RNAi *Ubr3* were expressed under the control of *Engrailed*-Gal4 driver visualised in red (*en*>mRFP). The yellow line delimits the anterior and the posterior regions. Rn pattern remains unchanged when *Ubr3* is depleted. (D) Model of Pri peptide functions during leg development: *pri* expression is spatio-temporally regulated during leg disc development. During L3 stage, Pri peptides activate Rotund and Spineless transcription factors, and EGFR and Notch signalling pathways. Therefore, Pri peptides coordinate transcriptional program and signalling pathways to ensure tarsal patterning. Then at larval-pupal transition, when *pri* is reactivated in the leg disc by ecdysone signalling, it induces Svb processing in a Ubr3 dependent manner. *Pri* expression is maintained during pupal stage in the leg disc. The module Pri/Ubr3/Svb is required for cell survival, but also morphogenesis and for maintaining epithelial integrity.

Our results reveal that Pri peptides functions in patterning during the larval stage are not mediated by Svb and are independent of Ubr3, suggesting the existence of additional Pri molecular targets. The ability of Pri peptides to control multiple cellular events through the activation of distinct factors within the same tissue over time illustrates their pleiotropic functions in the temporal control of development (Fig. 7D).

## Discussion

We took advantage of leg development features to identify novel putative smORF peptides and carried out a functional screen. We have thus shown that the family of smORF peptides, *i*.*e*. under 100 amino acid length, represents a reservoir of novel cellular and developmental regulators. Indeed, smORF peptides have been largely overlooked in genome annotations due to their small size and have remained underinvestigated so far. Focusing on leg development, we found that the most differentially expressed gene that encodes for smORF peptides controlling leg development is the *pri*/*tal* gene. Addressing Pri smORF peptide functions during leg development allowed us to dissociate larval and pupal roles, and thus to better understand its molecular action. Through their pleiotropic functions, Pri peptides by interacting with different actors trigger distinct molecular events, which synchronise tarsal patterning and morphogenesis required for harmonious leg development over time.

### smORF peptides provide a pool of novel developmental actors

Bioinformatics analyses have identified hundreds of putative smORF peptides, which have been classified in function of their origin. We focused on genes encoding smORF peptides that have never been studied because they were recently annotated (sCDS) or classified as lncRNA or pseudogenes. Importantly, genetic tools, allowing us to deplete their function, are available for half of these genes in stock centres for the fly community, and their use allowed us to identify a significant number of potential candidates for controlling development. Interestingly, 40% of smORF peptides display a motif, that could be useful to address its biological function. More than 10% of smORF peptides appear to be addressed to mitochondria, suggesting a tendency of the smORF peptides to localise to this organelle compared to the whole proteome (6%) as previously observed (Bosch et al., 2022; van Heesch et al., 2019). Furthermore, the phenotypes obtained are diverse, affecting cell survival, segment fusion or tissue growth, suggesting that smORF peptides are involved in all cellular processes. Indeed, recent studies showed that they can exhibit multiple subcellular localization (Na et al., 2022), with role in, for example, the regulation of calcium flux, the inhibition of protein activity (Jayaram et al., 2021), the antigen presentation (Niu et al., 2020) or the biogenesis of the respiratory chain (Liang et al., 2022). Obviously, further studies are now needed to better understand the function of these smORFs. Interestingly, we found that half of these putative regulatory smORF peptides have orthologs in vertebrates (Table S1), for instance, CG33169 (55AA) is encoding for the ortholog of human SMCO4 (59AA), a peptide of unknown function containing a transmembrane domain. Thus, *Drosophila* is a good model for identifying among the hundreds of existing smORF peptides new regulators of important cellular processes conserved in eukaryotes.

It is generally accepted that eukaryotic genes are monocistronic, *i*.*e*. they contain a single ORF. However, we found that a high proportion of predicted smORF peptides are located in the 5’UTR and 3’UTR, supporting the existence of polycistronic eukaryotic genes. Indeed, recent studies in *Drosophila* or vertebrates using mass spectrometry or ribosome profiling have shown that smORF peptides are indeed translated from the 5’ or 3’ UTR, even within the main ORF, demonstrating that polycistronic genes in eukaryotes are more widespread than expected (Chen et al., 2020; Fabre et al., 2022; Martinez et al., 2023). This highlights the potential of RNAs to code for several proteins, giving the possibility of greatly increasing the eukaryotic proteome. The challenge now is to define criteria or experimental approaches to select, among the thousands of smORF peptides, those most likely to have important regulatory functions in development.

Finally, our approach to search for putative smORF regulatory peptides, based on bioinformatic analyses of RNAseq data, is thus handable to *Drosophila* or any other organism. The use of criteria such as differentially expressed genes, expression level, or specific physiological conditions, as in our case before and after the ecdysone peak, can restrict the analyses to a smaller pool of genes. This may also increase the chances of finding a regulatory smORF peptide with specific spatio-temporal expression, which could help identify function or potential interactors. Here, these criteria highlighted the *pri*/*tal* gene, already known to be crucial for leg development (Galindo et al., 2007), thus validating our approach. As two peaks of *pri* expression occur during the development of the leg disc, we wondered whether they act with the same partners.

### Pri/Svb/Ubr3 module ensures leg tissue integrity during pupal stage

Pri peptides molecular function in epidermal differentiation, specifically during trichome formation, is mediated by Svb transcription factor and E3 ubiquitin ligase Ubr3 (Zanet et al., 2015). Here we demonstrated that the Pri/Svb/Ubr3 module is reused during leg metamorphosis, triggered by ecdysone signalling.

As previously shown in digestive stem cells, this module is required for protecting stem cells from apoptosis (Al Hayek et al., 2021; Bohère et al., 2018). Loss of function of one of these partners induces apoptosis during pupal stage and may be as a consequence, dramatic alteration of part of the tissue. As we showed that Pri peptides increase Ubr3/DIAP1 interaction in S2 cells, and that Ubr3 interacts with DIAP1 to protect cells from death (Huang et al., 2014), the couple Pri/Ubr3 could counteract apoptosis via promoting stabilisation and protection of DIAP1 *in vivo*. Even though Svb is also required for protecting cells from death, we did not find any effect of Svb on *DIAP1* expression as previously described (Bohère et al., 2018). However, Svb may be required later for tarsal growth and morphogenesis, as *svb* depletion during the pupal stage induces defects in leg development. Moreover, Svb is also required for epidermal differentiation and trichome formation not only in the tarsus, but also in the tibia (Kittelmann et al., 2018), and for joint formation (Pueyo and Couso, 2011). However, during the first hours of pupal leg development, we did not observe a role for Svb on Notch signalling or joint patterning as previously described (Pueyo and Couso, 2011). It might be due to the use of different genetic tools, like the *svb*::*GFP* knock-In line, which recapitulates the endogenous pattern of Svb.

The module Pri/Ubr3/Svb is operating at different stages of development and in different tissues in *Drosophila*, for instance in embryonic epidermis and in pupal leg, in which *pri* expression is temporally regulated by the ecdysone signalling (Fig. 7D) (Chanut-Delalande et al., 2014; Dib et al., 2021). Interestingly, the module Pri/Svb/Ubr3 is conserved among arthropods and regulates embryonic patterning (Ray et al., 2019), thus suggesting this module might be reiterated across arthropods development in different tissues and organs.

### Pri peptides synchronise signalling pathways and transcriptional program to ensure tarsal patterning

During the larval stage, we showed that the Pri/Svb/Ubr3 module is not mediating functions of Pri peptides for tarsal patterning. At midL3 stage, Pri peptides were known to control *spineless* (*ss*) and *rotund* (*rn*) expression (Natori et al., 2012; Pueyo and Couso, 2008), and we demonstrated that it is not mediated by Svb and Ubr3. Nevertheless, depletion of *Ubr3*, even at the larval stage, can induce apoptosis, which seems to alter epithelial leg disc morphogenesis.

Besides a fundamental role in initiating the tarsal transcriptional program, we found that Pri peptides are also required to activate EGFR and Notch signalling pathways. Notch activation correlates with tarsal sub regionalisation and segment emergence, which occurs in a Ubr3 and Svb independent manner. Our results suggest that *pri* is required for the second wave of EGFR activation in the tarsus, not in the initial EOC in the pretarsus, by regulating directly or indirectly the expression of *rhomboid* in concentric circles. Recently, it was shown in *Drosophila* embryonic tracheae that *pri* is also required for EGFR pathway activation in dorsal branches, thus supporting our data (Taira et al., 2021). Nevertheless, we observed that ectopic activation of the EGFR pathway is insufficient to mediate Pri functions, while EGFR seems to be required for patterning at the same time as *pri*.

Ss requires *pri* for inducing *rn* expression in the leg disc (Natori et al., 2012). Our experiments reveal that *rn*, as well as *ss*, was not able to restore tarsal patterning in the absence of *pri* and that *rn* is required for Notch signalling, thus suggesting that Pri peptides interfere several times in the regulatory cascade. Thus, we propose that Pri peptides are required concomitantly or reiteratively to activate key players, the EGFR and Notch signalling pathways and the transcription factors Ss and Rn, to synchronise the molecular events governing tarsal formation. Furthermore, cell non-autonomous properties of Pri peptides, allowing them to induce effects at a distance of several cell diameters from their domain of production, enable to activate Pri targets at comparable levels well beyond its expression domain. Pri’s cell non-autonomous effect is different from a gradient because Pri acts here as an interswitch: it either activates or does not activate its targets.

During the larval stage, Pri peptides are acting upstream of the signalling pathways and transcriptional cascade which govern tarsal patterning. Functions of Pri peptides are not mediated by Svb and Ubr3, thus suggesting that additional Pri peptides targets, direct molecular partners and/or indirect targets may exist. We speculate that the transcription factors Dll and Sp1 could interact with Pri peptides to mediate their functions as they control the same targets (Córdoba et al., 2016; Newcomb et al., 2018; Ruiz-Losada et al., 2018). However, deciphering the nature of their interaction will be a long-term effort as the interdependence of the key players and non-cell autonomous properties of Pri peptides renders difficult *in vivo* genetic approaches and interpretation of the effects of their manipulation.

Pri peptides may synchronise molecular events that are interdependent thanks to their non-cell autonomous properties. Thus, they couple signalling pathways and transcription program to orchestrate harmonious leg development. This plethora of molecular events regulated by Pri peptides is enabled by their pleiotropy. Indeed, they can simultaneously regulate different targets within the same tissue or even within the same cells. This pleiotropy is enhanced by their spatio-temporal transcriptional regulation, which relies on multiple enhancers (Dib et al., 2021). For example, the pulsatile expression of *pri* in the imaginal leg disc depends on several enhancers in the larval and pupal stages, of which only the pupal enhancers are regulated by ecdysone (Dib et al., 2021). We propose that Pri peptides rhythm *Drosophila* development by coordinating multiple and distinct cellular processes in space and time.

### Concluding remarks

The smORF peptides now represent a set of regulatory molecules capable of controlling cellular processes involved for example in development, metabolism, immunity and pathology. Furthermore, the example of the Pri smORF peptides illustrates the ability of a single peptide to induce a plethora of effects in a spatio-temporal manner by regulating distinct actors. Currently, thousands of smORF peptides have been shown to be actively translated, revealing the incredible coding potential of our genome as a source of novel bioactive molecules.

## Material and methods

### Bioinformatic smORF peptides prediction in imaginal leg discs

Imaginal leg discs were dissected in cold PBS 1X on ice from wandering L3 larvae, before the ecdysone peak, visualised by pre-spiracle eversion, and 2 hours After Pupal Formation (APF). Total RNA was extracted with Trizol® reagent (Ambion) according to the manufacturer’s protocol. Construction of RNA polyA+ bank and sequencing using paired-end 100bp reads were performed by IntegraGen®. Kallisto (Bray et al., 2016) was used for pseudo alignement of reads to a reference combining the Ensembl 74 annotations and additional lincRNAs from modENCODE (Brown et al., 2014; Young et al., 2012). We then used Sleuth for differential expression analysis (Pimentel et al., 2017). Small ORFs were predicted as in Mackowiak *et al*. using the transcriptomes generated in this study. The data discussed in this article have been deposited in NCBI’s Gene Expression Omnibus and are accessible through GEO series accession number GSE225561.

### Functional screen

Loss of function were induced by crossing *Dll*^EM212-Gal4^; tubGal80ts flies with lines expressing RNAi or gRNA and Cas9 under the control of UAS promoter at 29°C. Fly lines are available in Bloomington and VDRC stock centres, or were generated for this study. UAS-RNAi-*white* was used as the control. The lines giving a phenotype are listed in figure S1.

### Fly stocks

The *Drosophila* lines used in this study are *tal*^1^, *tal*^S18.1^, UAS-RNAi *pri* (Galindo et al, 2007), *Ubr3*^B^ (Zanet et al., 2015), *svb*^PL107^ (Delon et al., 2003), *Pri*I-Gal4 (generous gift from H Chanut-Delalande), UAS-*pri* (Kondo et al., 2007), *wingless*-Gal4, *dysfusion*640-LacZ (Córdoba and Estella, 2014). MARCM clones were generated by using the following fly line: *y, w, hs*-FLP, *tub*-Gal80, FRT19A; UAS::mcd8-GFP; *tub*-Gal4/TM6B, Tb (Bohère et al., 2018). The *Minute* clones in tarsus were generated by using the following line: yw; *Dll*, UAS-FLP; FRT82B, *Rps*, Ubi-GFP/Cyo-TM6b.

The following lines were available from Bloomington and VDRC stock centres: *Engrailed*-Gal4, UAS-mRFP (BL30557), UAS-RNAi *luciferase* (BL31603) and UAS-RNAi *white* (BL28980), both used as controls, UAS-RNAi *svb* (v41584), UAS-RNAi *Ubr3* (v22901, v106993).

The Knock-In of GFP protein at the C-terminal position of the Ovo/Shavenbaby protein in the endogenous locus was generated by CRISPR/Cas9 by InDroso® compagny.

### Hybridisation in situ

SmiFISH was performed as previously described (Couturier et al., 2019) and FLAP-X sequence was used to generate fluorescent probes. Larvae were dissected and fixed in PFA 4% in 25 min at room temperature, then washed in PBT (0,1% Triton100X) and permeabilized 20 min in PBT (0,5% Triton100X). Samples were washed in the wash buffer (4M urea in SSC 2X) and incubated with hybridization mix (4 M Urea, 8μL de SSC20X, 40μL of Dextrane 20%, 3,5 μL of Vanadyl complex at 10 mM, 1,5 μL of competitor DNA, 2,5μL of smiFISH probe and 1,5 μL of water) at 37° overnight protected from light. Samples were rinsed in the wash buffer and in SSC2X. Then, samples were washed in PBT (0,1% Triton100X) and leg discs were dissected and mounted in vectashield® medium (Vector Laboratory).

For rhomboid *in situ* hybridisation, probes (sense and the anti-sense) for *rhomboid* were synthesised according to standard procedures from LD06131 plasmid. Briefly, larvae were dissected in order to keep discs in PBS1X and fixed 20 min in PFA 4% at room temperature. Samples were washed in PBT (PBS1X/0,1% TritonX100), blocked 30 minutes in PBT (0,3% Triton X) and washed in PBT (0,1% Triton). Samples are permeabilized in Methanol/DMSO (90% /10%). Samples are rehydrated progressively and incubated overnight at 65°C with the denatured probe in the hybridisation buffer. Samples are washed, rehydrated progressively, incubated with anti-DIG (1/2000). Probe is then revealed with NBT/BCIP (Promega). Leg discs were dissected and mounted in a mix PBS/Glycerol.

### Immunofluorescence

Larval and pupal imaginal discs were dissected in PBS1X and fixed in PFA 4% during 25 minutes at room temperature, then washed in PBS1X. Samples are blocked in PBS/BSA 0,3%/Triton 0,3% during 1 hour. Primary antibodies are incubated overnight at 4°C. Then, samples were washed and incubated with secondary antibodies for two hours at room temperature, then rinsed in PBS1X and mounted in Vectashield® medium (Vector Laboratory). Antibody against Rotund (Rn) was obtained by immunizing guinea-pigs with the Roe (Roughened eye) full length isoform encoded by the *rotund* gene and sharing its last 450 C-terminal residues (including five Zn fingers) with the Rn isoform (St Pierre et al., 2002).

Contrary to Rn, *roe* is not expressed and has no function in leg tissues (St Pierre et al., 2002). GST-fused Roe was produced in E coli from a pGEX-Roe plasmid (del Alamo and Mlodzik, 2008), purified through a glutathione column and used to immunize the guinea pigs. Crude serum was used at 1:500. The antibodies used in this study are: anti-Spineless, generously given by J Yuh Nung (1/1000); anti-Distal-less generously given by R Mann (1/500), anti-GFP (Mouse, Roche) (1/500); anti-GFP (Rabbit, Torrey Pines) (1/500); anti P-ERK (P-p44/42 MAPK, Cell Signalling Technology), (1/200); 1/50, anti-Wingless (DSHB 4D4-s), (1/50); anti-Delta (DSHB C594.9B), (1/500); anti-Dcp-1 (Cell Signalling Technology), (1/200); anti-1S, (1/1500) (Zanet et al., 2015), anti-Deadpan (Abcam), (1/100). The secondary antibodies were coupled to Alexa Fluor 555, 488 or 647 (Invitrogen). The DNA was marked either with TO-PRO™-3 Iodide (642/661) or DAPI (Thermofisher).

### Co-immunoprecipitation and western blotting

S2 cells were transfected with pAc-V5::Ubr3, pAc-DIAP::GFP and pMT-pri. *Pri* expression was induced by CuSO4 at 1mM for 2 hours. Co-immunoprecipitation and western blotting were done as previously described (Zanet et al., 2015).

### Image acquisition

Experiments with fluorescent markers were obtained using microscope Leica sp8. Experiments requiring white light like in situ hybridization or adult leg observation are acquired with microscope Nikon Eclipse 90i.

## Acknowledgments

We thank Jan Yuh Nung, Richard Mann, Carlos Estella and Hélène Chanut-Delalande for sharing antibodies and fly lines. We also thank the Developmental Studies Hybridoma Bank for antibodies, the Bloomington *Drosophila* Stock and Vienna *Drosophila* Ressource Centres for fly strains. We thank LITC platform for imaging (https://www-litc.biotoul.fr/). We thank Magali Suzanne for critical reading of the manuscript. We thank Hélène Chanut-Delalande and Cédric Polesello for helpful comments, and Alexia Rivero for helping visualising sequencing data.

