## Supplementary_Information for "The pleiotropic functions of Pri smORF peptides synchronise leg development regulators"

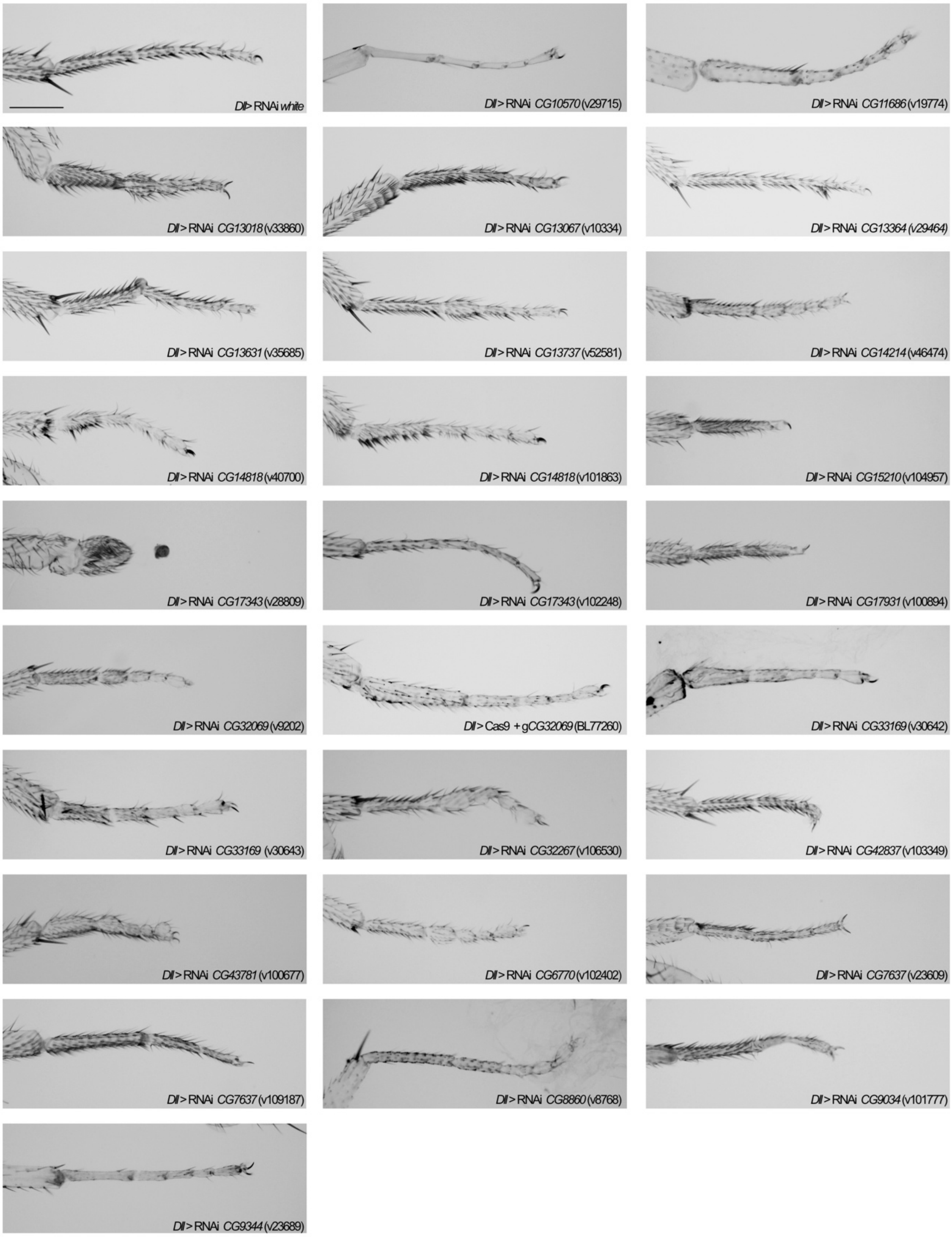


**Figure S1**

**Figure S1 : Loss of function of smORF peptide in tarsus induces multiple developmental defects**

Here are shown the different phenotypes and defects obtained following depletion of smORF peptide encoding genes identified in the functional screen. Loss of function was induced by expressing UAS-RNAi, or UAS-gRNA and UAS-Cas9, under the control of *Dll*-Gal4 driver. RNAi lines used are specified on each picture with the name of the CG targeted. We observed abnormal fusion of tarsal segments, defects in tarsus growth and cuticle formation, showing that smORF peptides identified here control different cellular processes. Scale bar = 200µm.

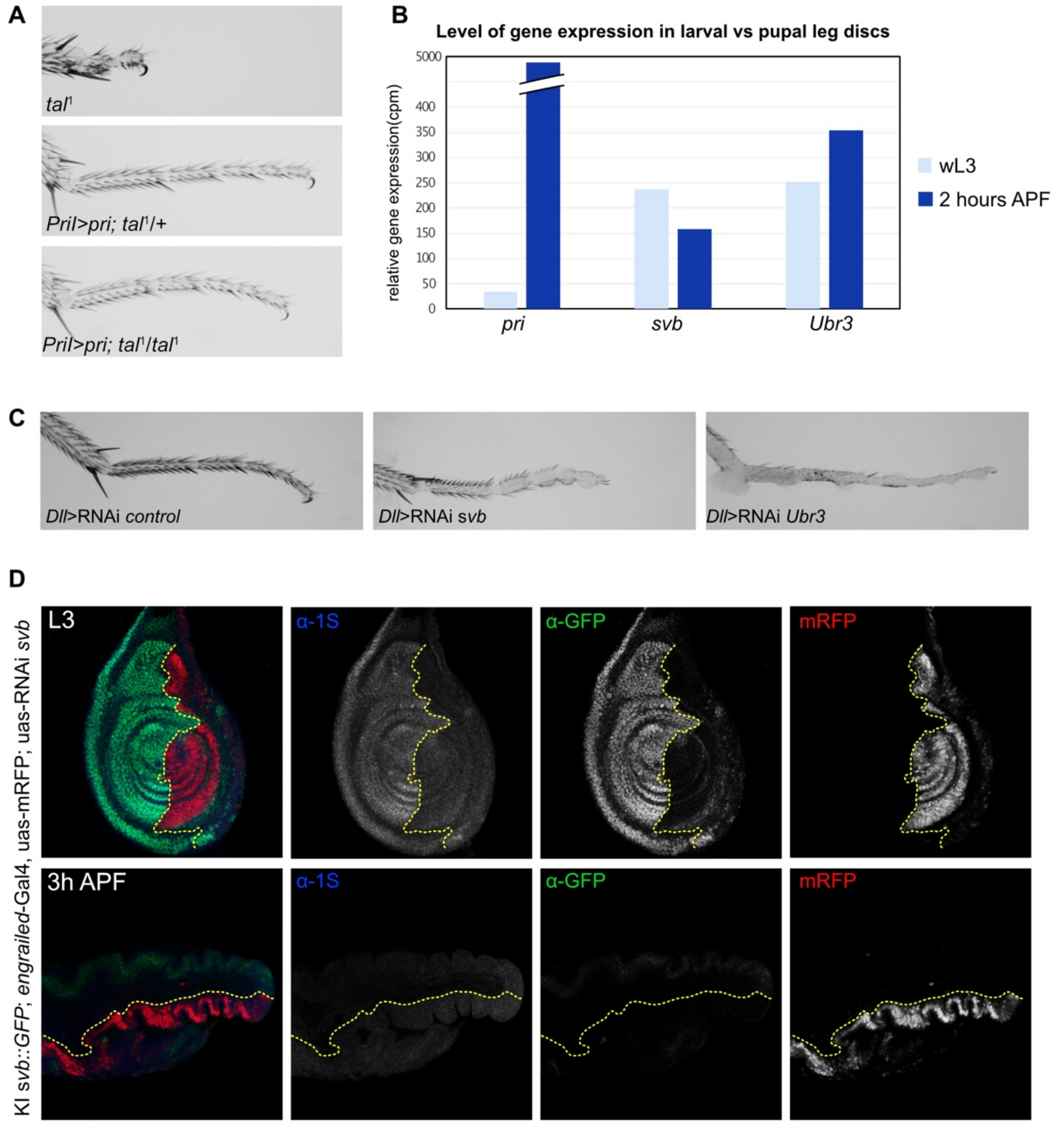


**Figure S2**

**Figure S2 : Shavenbaby (Svb) and Ubr3 are required for leg morphogenesis**
(A) Morphology of the tarsus of *tal*[1] mutant is rescued when *pri* is ectopically expressed under the control of *Pril*-Gal4 driver at 18°C. (B) RNA-seq analysis on imaginal leg discs at wandering L3 stage (wL3) and at pupal stage 2 hours APF (After Pupal Formation) show a massive up-regulation of *pri* expression, whereas *svb* and *Ubr3* expressions remain stable. (C) Depletion of *svb* or *Ubr3* in the tarsus, induced by the expression of RNAi under the control of *Dll*$^{EM212}$-Gal4 driver, alters morphogenesis. In the absence of *svb*, the segments of the tarsus are shorter and the trichome pattern is affected. Loss of function of *Ubr3* leads to a dramatic phenotype, the integrity of the epithelium and cuticle differentiation are impaired. (D) Expression of UAS-RNAi *svb* in the posterior domain (*En*-Gal4) of the leg disc in KI *svb*::*GFP*, marked with the mRFP, demonstrates that endogenous Svb protein is fused with the GFP and localises ubiquitously within the leg disc. Anti-1S staining shows that Svb is under the full length repressor form. At the larval-pupal transition, Svb is processed, and remains under the short activator form during pupal leg development.

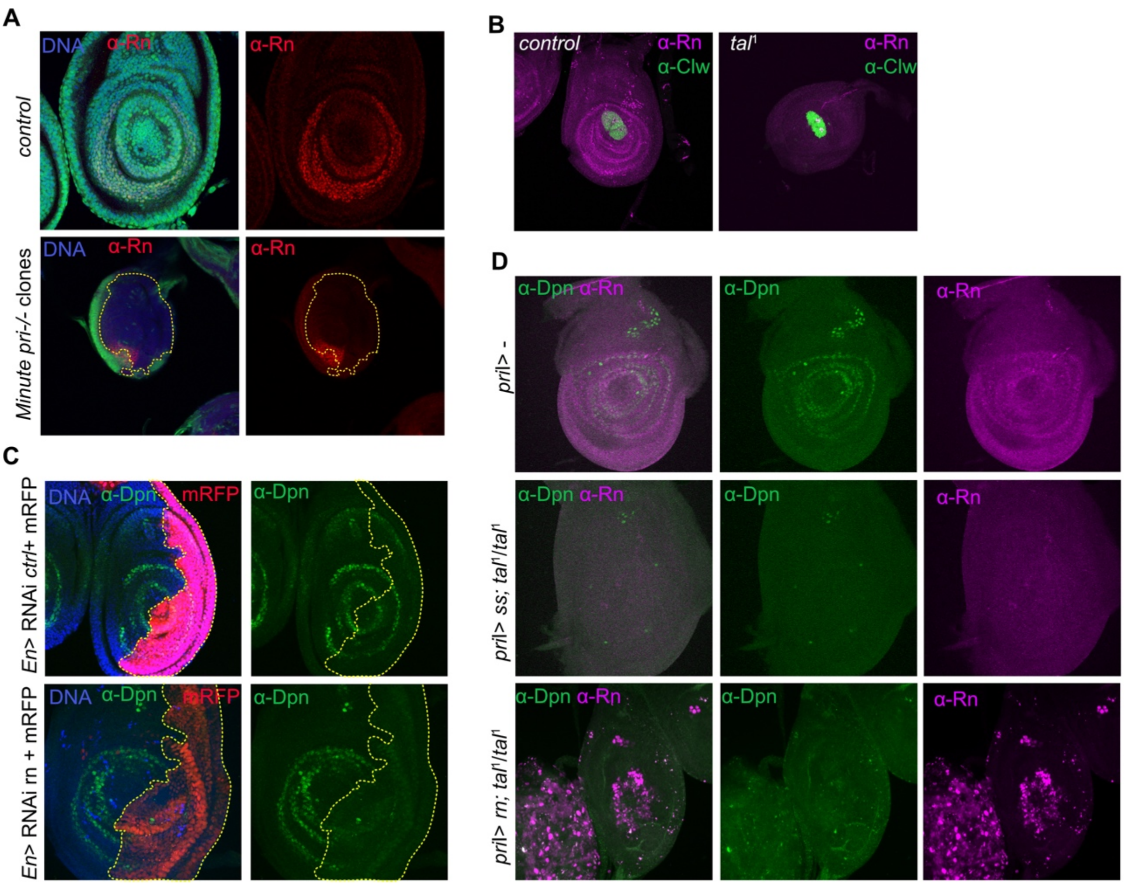


**Figure S3**

**Figure S3 : Interdependence between Pri peptides, Notch signalling and tarsal transcription factors**

(A) Rn immunostaining in *pri*$^{-/-}$ (*tal*$^{S18}$) clones induced in the *Minute* cellular context. Clones are indicated by the absence of GFP. The control displays no clone and Rn protein is localised in the presumptive region of the tarsus. The *pri*$^{-/-}$ clone is large enough (outlined by the yellow dashed-line) to encompass most of the leg disc, Rn pattern is dramatically affected. Note that Rn is activated beyond the GFP positive zone, in cells that are not expressing *pri*, showing Rn activation in cell non-autonomous manner. (B) Anti-Clawless (Clw) (1/200; Kojima et al, 2005) staining is specific from the pretarsus and is present in *tal*$^{1}$ mutant, showing that Pri peptides are not required for pretarsus patterning. (C) *Rotund* (*rn*) was depleted by RNAi (BL65347) specifically in the posterior region of the disc under the control of *Engrailed*-Gal4 (*En*) driver. The RNAi control (ctrl) used here is RNAi *white*. Anti-Dpn staining is absent when *rn* is deleted, showing that Rn is required for activating Notch signalling pathway. (D) Rescue experiments have been conducted by expressing ectopically *spineless* (*ss*) (BL78354) or *rotund* (*rn*) (BL7404) in *tal*$^{1}$ mutant background under the *pri*I-Gal4 driver, *i.e.* in the presumptive tarsal region. We observe that neither Ss nor Rn is sufficient to activate Notch signalling in the absence of *pri* since anti-Dpn staining is missing.

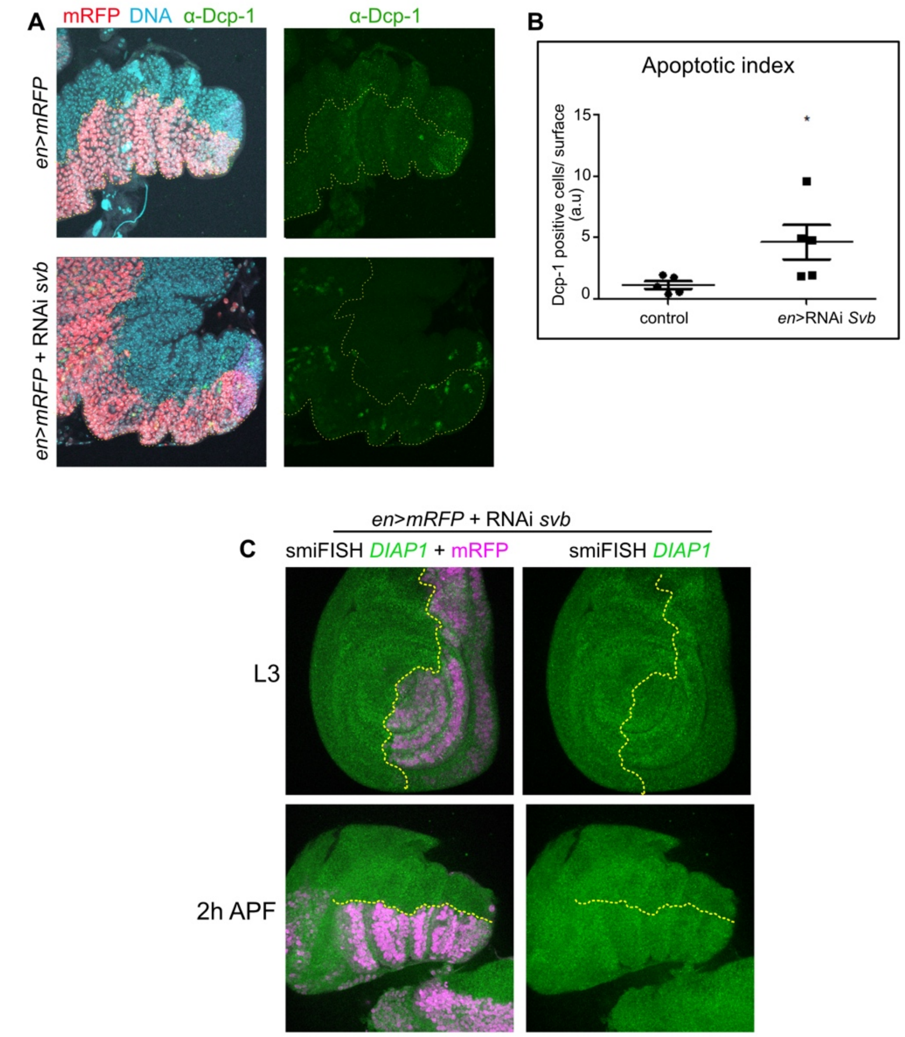


**Figure S4**

**Figure S4 : Svb is required for preventing cells to enter in apoptosis**
(A) *svb* was specifically deleted in the posterior *engrailed* domain (*en*-Gal4; UAS-RNAi *svb*, outlined by the yellow dashed-line) and apoptotic cells were stained with anti-Dcp-1 antibody (in green). We observed an increase in Dcp-1 positive cells in the *engrailed* domain compared to the control domain. (B) Apoptotic index reflects the number of apoptotic cells between the posterior (*engrailed*) and the anterior domains of the tarsus. It is the ratio between the number of apoptotic cells between the posterior zone of the leg, where the RNAi *white* (control) or the RNAi *svb* are expressed under the control of *en*-Gal4 driver, and the anterior region. Dcp-1 intensity signal was measured with ImageJ. Depletion of *svb* induces an increase of apoptotic cells. n=5 for each genotype. (C) Fluorescent *in situ* hybridization in imaginal leg discs of *DIAP1* mRNA in L3 and 2 hours APF. RNAi *svb* was expressed under the control of *engrailed*-Gal4 driver (*en*>) in posterior region, visualised with mRFP (purple).No significant change in *DIAP1* mRNA level is observed.

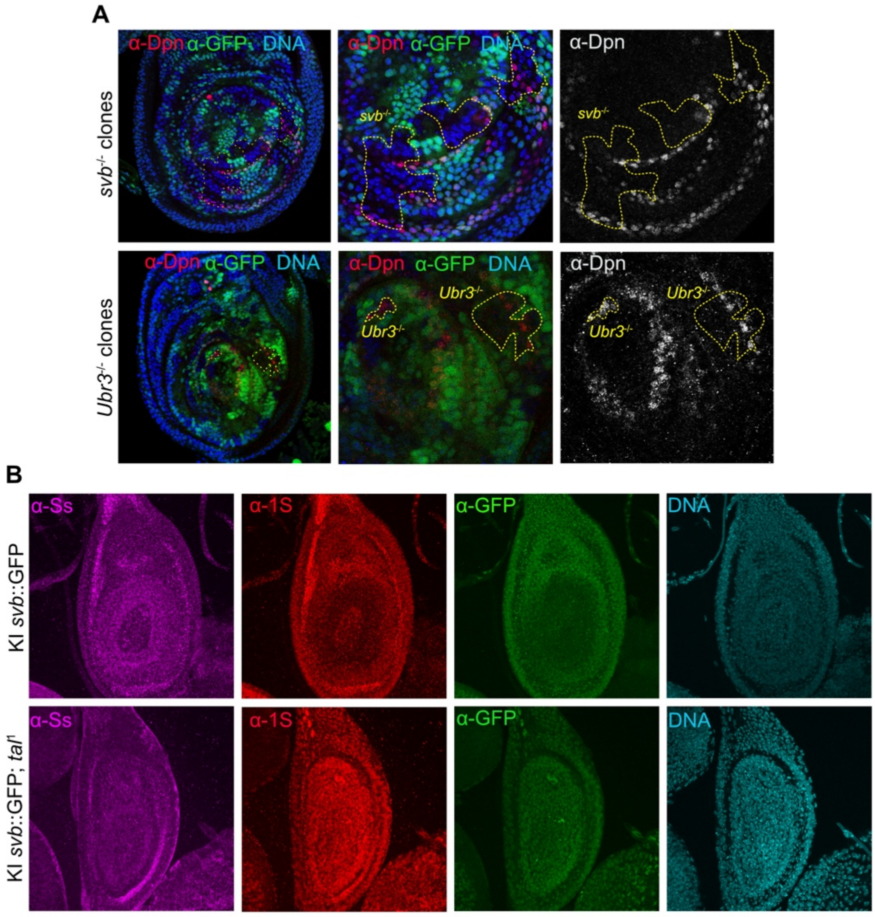


**Figure S5**

**Figure S5 : Svb and Ubr3 do not regulate Notch signalling pathway at larval stage**
(A) *svb*$^{-/-}$ (*svb*$^{PL107}$) and *Ubr3*$^{-/-}$ (*Ubr3*$^{B}$) clones were generated in leg discs, that were stained with anti-GFP and anti-Dpn antibodies to visualize the activity of Notch signalling pathway. The clones are GFP negative. The absence of *svb* or *Ubr3* does not affect Notch signalling, as Dpn staining is present in clones. (B) Anti-1S and anti-GFP stainings in KI *svb*::*GFP* and in *tal*$^{1}$ mutant background reveal that Svb is fully degraded at midL3 in the tarsal presumptive region, marked here with the anti-Spineless (Ss) antibody. In *tal*$^{1}$ mutant background, Svb full degradation does not occur, showing that *pri* is required in this process.

**TableS1**

Here is the list of small ORF that were bioinformatically predicted. Genomic position, name and length (in AA) are shown. Predicted motifs (signal peptide, transmembrane domain, mitochondrial targeting sequence) are also specified. The phyloCSF score is reflecting the conservation of the ORF between the 12 *Drosophila* species, and is considered to be relevant above 50. The type refers to the position of smORF. The annotated smORF, which have a name, were either functionally studied, or annotated based on the conservation of protein sequence between eucaryotes. Most CG number are smORF that were recently annotated. Pseudogene means the gene is considered as non functionnal. Non-coding means that the smORF is localised in non-coding RNA. CDS means the smORF in localised within the coding sequence of a canonical gene, UTR5 upstream and UTR3 downstream of the coding sequence. Other means the smORF is localised in intron or in intergenic region. Differential expression analyses were done between RNAseq data from larval and pupal leg discs. Qval is the p-value of the statistical test, logFC is the log2fold change of expression level between the two conditions, TPM-L, transcripts per million in larval discs, TPM-P, transcripts per million in pupal discs.
